# Mathematical modeling of Erk activity waves in regenerating zebrafish scales

**DOI:** 10.1101/2021.01.25.428093

**Authors:** L.D. Hayden, K.D. Poss, A. De Simone, S. Di Talia

## Abstract

Erk signaling regulates cellular decisions in many biological contexts. Recently, we have reported a series of Erk activity traveling waves that coordinate regeneration of osteoblast tissue in zebrafish scales. These waves originate from a central source region, propagate as expanding rings, and impart cell growth, thus controlling tissue morphogenesis. Here, we present a minimal reaction-diffusion model for Erk activity waves. The model considers three components: Erk, a diffusible Erk-activator, and an Erk-inhibitor. Erk stimulates both its activator and inhibitor, forming a positive and negative feedback loop, respectively. Our model shows that this system can be excitable and propagate Erk activity waves. Waves originate from a pulsatile source which is modeled by adding a localized basal production of the activator that switches the source region from an excitable to an oscillatory state. As Erk activity periodically rises in the source, it can trigger an excitable wave which travels across the entire tissue. Analysis of the model finds that positive feedback controls the properties of the traveling wavefront and that negative feedback controls the duration of Erk activity peak and the period of Erk activity waves. The geometrical properties of the waves facilitate constraints on the effective diffusivity of the activator, indicating that waves are an efficient mechanism to transfer growth factor signaling rapidly across a large tissue.

**Significance statement:** Signaling waves represent a possible mechanism of spatiotemporal organization of multicellular tissues. We have recently shown that waves of activity of the kinase Erk control osteoblast regeneration in adult zebrafish scales. Here, we present a detailed characterization of a mathematical model of these signaling waves. We show that a source region poised in an oscillatory state can broadcast traveling waves of Erk activity in the surrounding excitable tissue. The dynamics of the source control the number and frequency of waves. Geometrical arguments support the notion that excitable Erk waves are an effective mechanism to transport growth factor signaling across a large regenerating tissue.

## Introduction

The proper function of multicellular systems requires tight coordination of their cellular components. Coordination across tissues is often provided by non-uniformly distributed signaling molecules (1–3). In the classic “morphogen gradient” model, a certain information-carrying molecule, the morphogen, is distributed in a graded manner across the tissue, and cell behaviors are instructed differentially by morphogen local concentration or its dynamics (3–5). These morphogens can be transcription factors, as in the case of Bicoid in *Drosophila* embryos, or extracellular ligands that bind to a transmembrane receptor and activate signaling cascades. In a simple model of gradient formation, morphogen molecules are produced in a localized source, diffuse in the extracellular space, and are absorbed/processed (“degraded”) by cells in a characteristic timescale. Under these conditions, a gradient would form, possessing an exponential decay profile with a length scale equal to the square root of diffusivity divided by degradation rate. While morphogens gradients can form quickly in small tissues, it might be difficult to establish such diffusion gradients spanning large tissues, as the time needed to distribute molecules by diffusion scales with the square of the distance. Moreover, the time needed for morphogen concentration to approach its equilibrium value can vary significantly across the gradient (6), suggesting that for large tissue a static morphogen gradient might be difficult to generate and/or maintain. As a consequence, while several examples of morphogen gradients have been described in embryonic tissues of size of 10-100 microns, the role or mechanism of establishment of these gradients in larger tissues of order of a millimeter or longer remains poorly understood (7). Active transport, for example through specialized cytonemes (8, 9), can provide faster morphogen spreading. Alternatively, coupling diffusion with positive feedback can transport information across large spatial scales by generating waves (10–13). Waves can propagate quickly across a tissue and maintain their intensity, providing efficient information transport (14, 15). Thus, waves are an alternative mechanism to morphogen gradients for the regulation of cellular dynamics in large tissues.

Signaling waves might be particularly important in the control of adult tissue regeneration. Regeneration is the process through which tissues regain a functional form following injury. Tissue regeneration requires precise control of cellular dynamics across a wide range of temporal and spatial scales (16). Several of the pathways that are important during development are reactivated in regeneration (17), but it is unclear how these signals are coordinated across the large spatial dimensions of adult regenerating tissues. Extracellular signal-Regulated Kinase (Erk) is a signaling component which has been implicated in many developmental contexts, including regeneration (18–20). Several feedback systems confer Erk activity a variety of dynamic behaviors which can lead to different outcomes in a context-dependent manner (21, 22). Understanding how these feedback mechanisms are explored in different biological processes is likely to reveal important regulatory principles. In particular, it has been shown that Erk oscillations can be transmitted to nearby cells and propagate as waves, thereby coordinating cellular dynamics across multiple cells (23–26). For example, in response to a wound in the mouse skin, epithelial cells collectively migrate toward the wound (25, 27). This process is recapitulated in wound assays of MDCK cell cultures where it has been extensively studied. Cells most proximal to the wound first move toward the injury site, inducing stretching forces on the neighboring cells (24, 28–30). This mechanical deformation likely triggers activation of a disintegrin and metalloprotease 17 (ADAM17), which in turn drives the release of membrane-tethered epidermal growth factor (EGF), signaling to neighboring cells through the epidermal growth factor receptor (EGFR) (24). Erk activation through the EGFR signaling cascade induces cell contraction in follower cells. Contraction in these cells exerts stretching forces on the next follower cells, prompting another round of Erk activation. This positive feedback loop between mechanical forces and Erk activation (24) can result in traveling waves. It has been proposed that the coupling of forces and Erk activity facilitates long-range order and migration in the direction of the wound (31), whereas mechanical forces alone would tend to lose directionality and strength while spreading across a tissue (32, 33).

Recently, we have reported Erk activity waves *in vivo* in regenerating osteoblast tissue in zebrafish scales (34). These waves have properties of reaction-diffusion waves in an excitable medium. Here we present a detailed characterization of a mathematical model of Erk activity propagation in zebrafish scales. Our model shows that coupling a localized oscillatory source region with a surrounding excitable tissue can result in the generation of repeated excitable waves. Thus, our model suggests that tuning the dynamic properties of the source region is a simple strategy to control wave generation and, ultimately, tissue growth.

## Results

### A three-component excitable system generates Erk activity waves

Zebrafish scales are millimeter-size bone disks that form a protective skeletal array on the body of the fish (35). The scale bone is covered by a monolayer of bone matrix secreting osteoblasts. After scale loss, a new osteoblast pool regenerates and reforms the bone. The osteoblast tissue forms first by differentiation of an unknown progenitor, then osteoblasts proliferate and finally increase in size without cell division (cell hypertrophy) (34, 36, 37). The hypertrophic phase is coordinated by repeated waves of Erk activity, which can be visualized using a transgenic biosensor expressed specifically in osteoblasts (Fig 1A). Using image analysis methods, we were able to build dynamic heat maps of Erk activity in the entire osteoblast population of regenerating scales—see Figure 1A and (34). This approach revealed repeated Erk activity waves which originate from a central source region and propagate outward as expanding rings (Fig. 1A, (34)). These waves move at a speed on the order of 10 μm/h and cross the entire scale in a few days; Erk is activated in 3 h and de-activated in 5 h, thus generating a peak 50-100 μm wide (34). The dynamic properties of the central source region suggest that it could be a dynamic system operating in an oscillatory regime, while the rest of the scale would be excitable. An excitable system is a non-linear system that has a single stable point and displays small responses to small stimuli but undergoes a large excursion in the phasespace in response to perturbations above a certain threshold (38, 39). Multiple excitable units can be coupled by diffusion and generate waves. Compatible with this idea, we found that wave initiation and propagation depend on Fibroblast Growth Factor Receptor (Fgfr) signaling, which is activated by extracellular diffusible ligands of the Fibroblast Growth Factor family (Fgfs) (34). Thus, Fgfs are candidates for propagating Erk activity from one cell to the next. In order to form an excitable system that can sustain a wave, Erk active cells need to produce or stimulate Fgfs, thus generating a positive feedback loop. In addition, cells at high Erk activity switch off after a certain time, indicating that one or multiple Erk inhibitors are activated in Erk active cells, generating a negative feedback loop. Thus, the minimal components of the proposed model are: Erk itself, an Erk-activator, and an Erk-inhibitor. The activator can diffuse in the extracellular space, while the Erk kinase and its inhibitor are confined within each cell. We take a coarse-grained continuum approach and describe the extracellular movement of the Erk activator by simple diffusion (Fig. 1B). These assumptions lead to the following model for Erk activity:

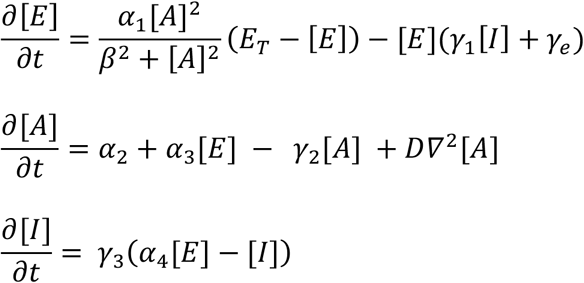

where [*E*], [*A*], and [*I*] represent the concentrations of active Erk, diffusible activator, and inhibitor, respectively. For simplicity, reaction terms are described according to the law of mass action with the exception of the rate of Erk activation by the activator, expressed as a Hill function of the activator concentration. A non-linear term in the positive feedback regulation between the activator and Erk is needed for the system to be excitable. While we chose to describe the dependency of Erk on activator levels as a non-linear function, similar results would be obtained by making different choices for the non-linearity. In the model, α_1_ is the Erk activation rate at saturating activator, *β* is the AC_50_ of Erk activation by the activator, *E_T_* is the mass-conserved sum of active and inactive Erk, *γ*_1_ is the Erk inactivation rate by the inhibitor *I, γ_e_* is the Erk autonomous deactivation rate, *α_2_* is the activator production rate, *α_3_* is the Erk-dependent activator production rate, *γ*_2_ is the autonomous activator decay rate, *D* is the diffusion constant of the activator, *γ*_3_*α*_4_ is the inhibitor production rate and *γ*_3_ is the inhibitor autonomous degradation rate. By nondimensionalizing the concentrations: [*E*], [*A*] and [*I*] by the total amount of Erk *E_T_* and expressing them without brackets, we obtain the equations:

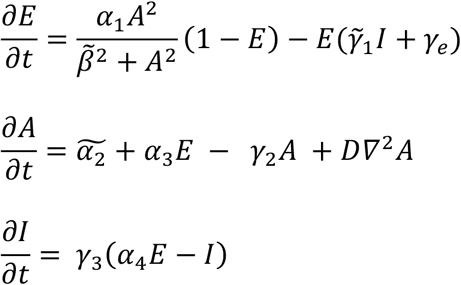

where 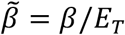, 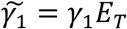, and 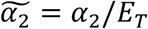 For sake of simplicity, we drop the tilde signs from 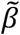, 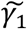, and 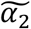 hereafter. In order to tune the system to an oscillatory state in the source region (Fig. 1C), instead of an excitable state, we introduced in that region a constant term of production of the activator *α_2_*. This term can be interpreted as a basal activator production by osteoblasts in the source region or a contribution from an external cellular pool. This term is null in the excitable region. Numerical solutions of the previous equations demonstrate that the system can generate traveling waves of Erk activity (Fig. 1D). As expected, linear stability analysis shows that, with the chosen set of parameters, one oscillatory fixed point exists in the source region (limit cycle, Fig. 1E), while outside the source region the system is excitable (Fig. 1F). Thus, oscillations of activator levels (and Erk activity) in the source trigger the propagation of excitable waves. The features of the excitable wave can be understood through the following arguments. The leading edge of the wave (inhibitor levels low) can be approximated with a bistable system, with stable fixed points at high and low Erk activity. Activator diffusion triggers excitations in the tissue in front of the leading edge. In that region, the system will leave the basin of attraction of the low Erk activity state and move toward the high Erk activity state. In turn, the accumulating active Erk produces its own inhibitor. As the inhibitor accumulates, the level of active Erk at the high stable fixed-point decreases until the system arrives at a saddle-node bifurcation, that results in only one stable point at low Erk activity, remaining (Fig. 1G). The system cannot be excited again until the inhibitor has substantially degraded (refractory period). Thus, our analysis points to a simple dynamical picture for Erk waves in scale regeneration.

**Fig. 1.**
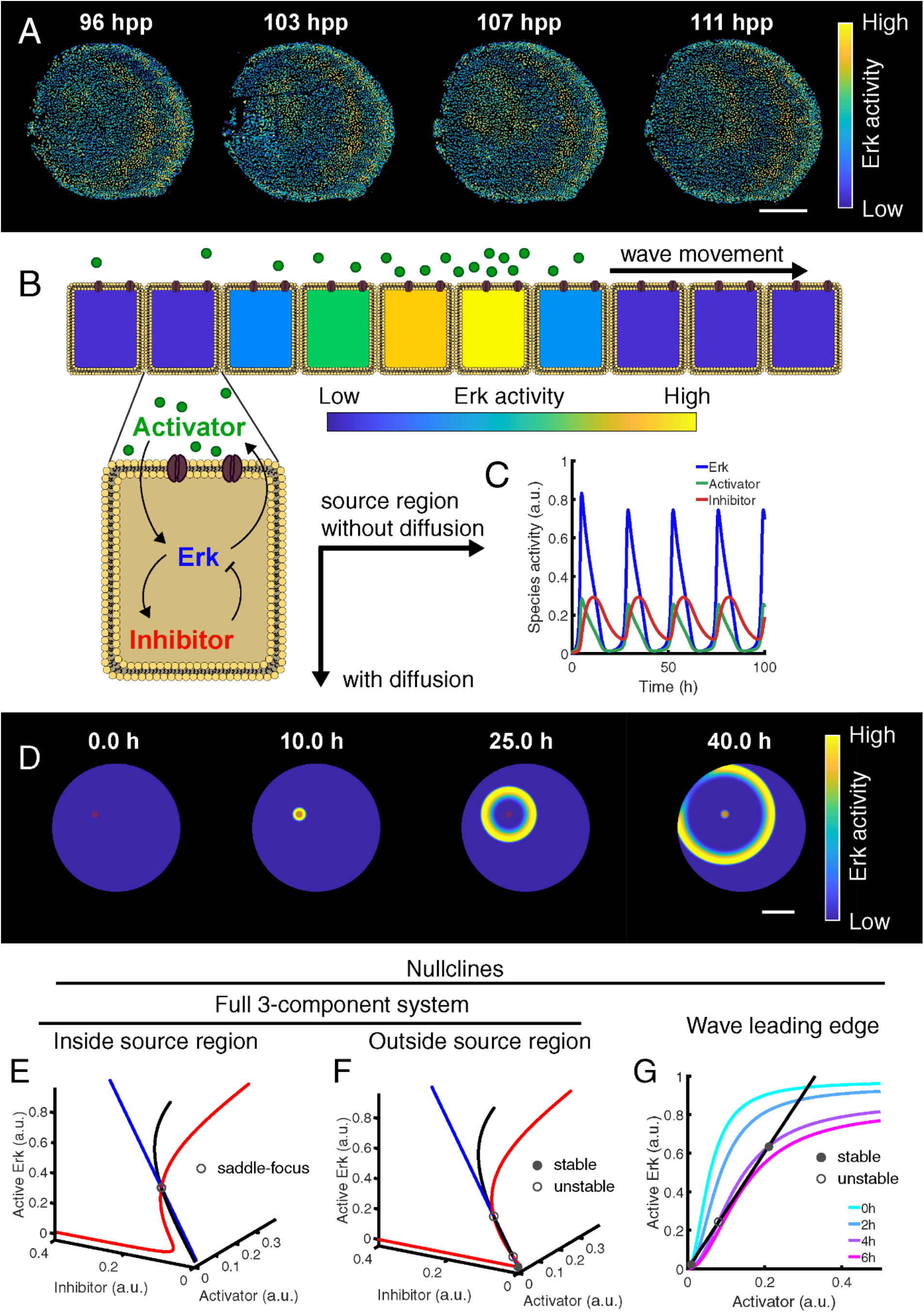
Model formulation and stability analysis. (A) Erk activity in regenerating scales *in vivo,* measured using an Erk Kinase Translocation Reporter (Erk KTR). The Erk KTR sensor provides a proxy of Erk activity through measurement of its subcellular localization. The sensor contains both a Nuclear Localization Signal and a Nuclear Exclusion Signal and preferentially localizes to the nucleus when unphosphorylated and to the cytoplasm when phosphorylated (40). Thus, a readout of Erk activity at cellular resolution is calculated by measuring the ratio of nuclear and cytoplasmic Erk KTR signals. The data shown was imaged and analyzed in (34). hpp: hours post plucking. (B) Schematics of theoretical model including a diffusible extracellular activator, Erk, and an inhibitor of Erk. Arrows between chemical species indicate feedback. (C) Numerical solution of the dynamical system in the source region and when diffusion is null. (D) Numerical solution of the reaction-diffusion dynamical system. (E, F) Stability analysis of the system when diffusion is null in (E) and outside (F) the source region. Curves indicate pairwise intersections between nullcline surfaces. (G) Leading edge fixed-point analysis: nullclines in the leading edge of the wave, where the inhibitor is approximated with a constant concentration. The activator nullcline (black) does not depend on inhibitor and therefore does not change over time, whereas the slope of the Erk nullcline varies (in color). Circles: fixed points at 4 h from simulation start (stability is indicated). Scale bar: 250 μm.

To understand which parameters control wave properties, namely wave frequency and speed, we performed a parameter sensitivity analysis by varying a single parameter at a time. The phenomenology of source oscillation and wave propagation can change while varying parameters. For example, for some parameter variations, damped waves originate from the source or the source/excitable region is locked in a high or low Erk activity state (Supplementary Figure 1). However, in our *in vivo* observations, each oscillation of the source corresponded to the generation of a wave reaching the edge of the tissue. Therefore, we limited our analysis to parameter values that generate a phenomenology that corresponds to these observations (see Methods for details). We found that waves with a well-defined speed and frequency are generated in a wide range of the parameter space with some parameters having broader ranges than others. As the parameters are varied outside this range, stable waves are replaced by either unstable dispersions which lose amplitude over time or a stable propagating front of high Erk activity, indicating the establishment of a bistable system (Fig. 2).

**Fig. 2.**
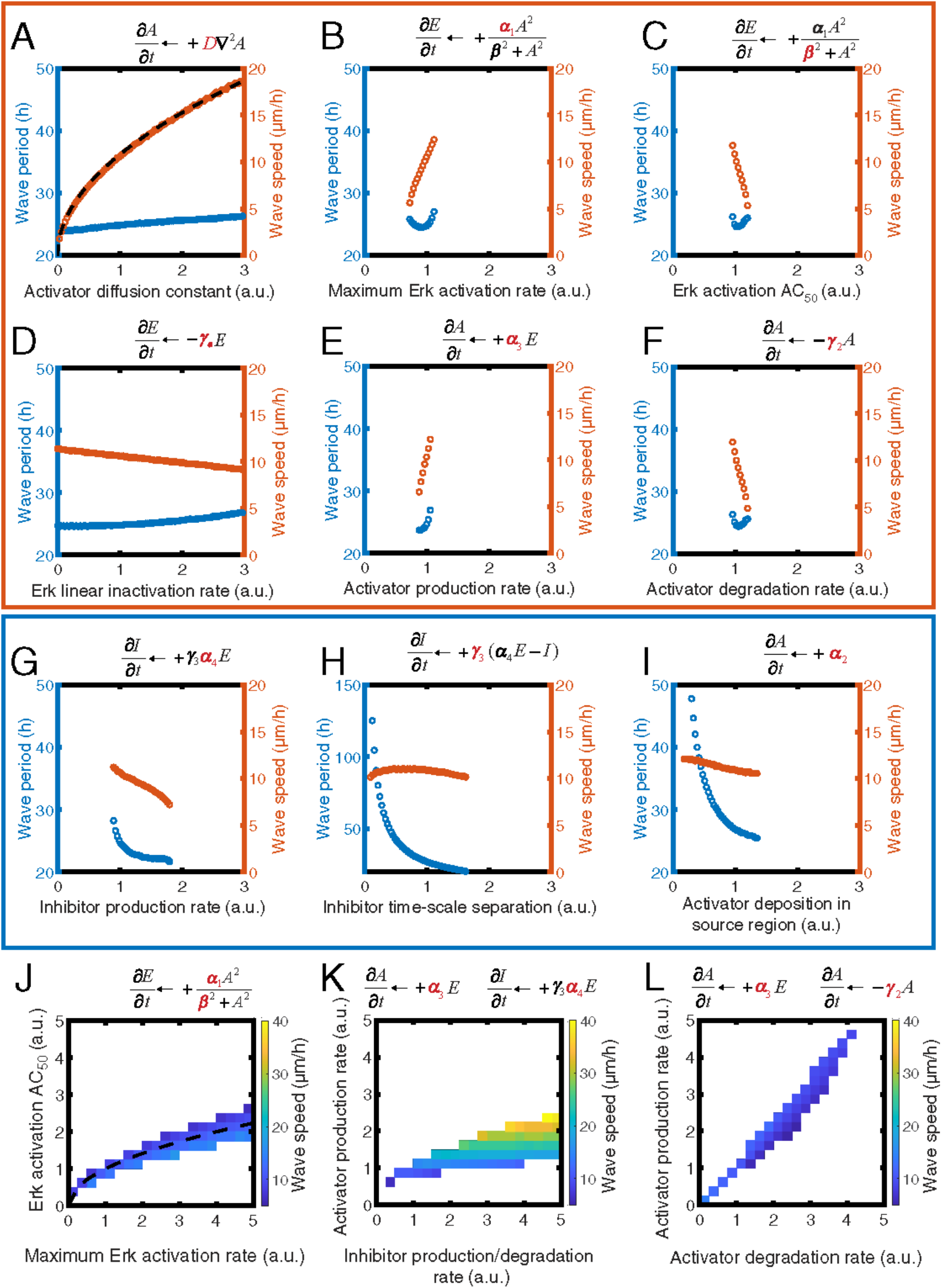
Parameter analysis and sensitivity. Wave speed and period varying the parameters of the model: D (A), *α*_1_ (B), *β* (C), *γ*_e_ (D), *α*_3_ (E), *γ*_2_ (F), *α*_4_ (G), *γ*_3_ (H), and *α*_2_ (I) individually and covarying *α*_1_/β (J), *α*_3_/*α*_4_ (K), and *α*_3_/*γ*_2_ (L). Parameters are expressed relative to the reference value in our standard simulation (see Table 1). Predicted square root dependencies in (A, J) are plotted as dashed black curves: (A) *υ* = *υ*_0_*D*^1/2^ where *υ*_0_ = 10.75 μm/h is velocity for the standard set of parameters and D is expressed in relative units and (J) square root dependency: 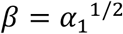. Wave speed and period are calculated for regions of parameter space which feature periodic stable waves. Orange box: parameters that affect mainly wave speed. Blue box: parameters that affect mainly wave period. Please note that *γ*_1_ and *α*_4_ together determine the action of the inhibitor by controlling its activity towards Erk and its levels, respectively. Thus, the dependencies of the wave period and speed on those parameters are identical, and the plot for *γ*_1_ was omitted.

**Table 1:** Parameters of the standard simulation. Note that to fully nondimensionalize the equations of the mathematical model, we would need to specify the concentration of total Erk [*E_T_*]. We lack experimental data on this parameter; however, experiments in different cell types argue for values in the range of [*E_T_*] ~0.1 −1 μM (49). Notice that these parameters are slightly different from those of (34), as we found that by rescaling all time-dependent parameters, we could tune *α*_2_ to obtain any wave frequency seen experimentally.

| Parameter | Value | Description |
| --- | --- | --- |
| $\alpha_1$ | 12.6 h<sup>-1</sup> | Maximum Erk activation rate |
| $\beta$ | 0.35 | Erk activation AC<sub>50</sub> |
| $\gamma_1$ | 15.4 h<sup>-1</sup> | Erk linear inactivation rate |
| $\gamma_e$ | 0.14 h<sup>-1</sup> | Erk linear inactivation rate |
| $\alpha_2$ | 0.112 h<sup>-1</sup> (inside source)<br>0 (outside source) | Activator deposition rate |
| $\alpha_3$ | 3.9 h<sup>-1</sup> | Activator production rate |
| $\gamma_2$ | 11.8 h<sup>-1</sup> | Activator degradation rate |
| $D$ | 566 $\mu\text{m}^2/\text{h}$ | Activator diffusion constant |
| $\gamma_3$ | 0.14 h<sup>-1</sup> | Inhibitor time-scale separation |
| $\alpha_4$ | 0.5 | Inhibitor production rate |

The theory of chemical waves predicts that the speed of the wave is determined by the activator diffusion constant *D* and by the timescale of activation *τ* via 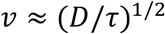. As predicted, the speed of Erk activity waves scales as the square root of the diffusion constant (Fig. 2A). Furthermore, our analysis demonstrates that parameters controlling the activator-Erk positive feedback loop have a larger impact on wave speed than its frequency (Fig. 2B-F). This can be intuitively understood by the fact that the positive feedback loop between Erk and its activator dominates in the leading edge. On the other hand, parameters controlling inhibitor dynamics mainly impact wave frequency as they control the duration of the refractory period and thus the intrinsic period of excitable excursion. An interesting exception is the rate of deposition of the activator at the source (*α*_2_), which controls the period of oscillation of the source. When the period imparted by *α_2_* is longer than the intrinsic oscillation period of the excitable region (Fig. 2G-H), this period imparted by *α_2_* will determine the tempo of wave generation. However, when the oscillation period at the source is shorter than the intrinsic oscillation period of the excitable system, this latter intrinsic oscillation period will limit the rate of wave generation (Fig. 2I).

Sensitivity analysis also indicates that several parameters have a similar impact on wave speed and frequency, and this co-dependency is strengthened by changing two parameters simultaneously (Fig. 2J-L). For some of these parameters, this co-dependency is intuitive. For example, the speed of the Erk wave is controlled by the dynamics of the leading edge at concentration *A≪β;* therefore, the term controlling Erk activation by the activator effectively reduces to *α*_1_,/*β*^2^, which effectively captures the co-dependency between the two parameters (Fig. 2J). This analysis suggests that while our model has several parameters that are not known experimentally, the emergent properties of the system belong to a few general scenarios. Furthermore, parameter values are constrained by the dynamic features of Erk oscillations and waves.

### The dynamic properties of the source region can impact how wave generation responds to perturbations

In the proposed model, the Erk activator accumulates in the source region, forcing it to an oscillatory regime (Fig. 3A). In an alternative model, external activator deposition may be oscillatory (Fig. 3B). In principle, these two different scenarios could be distinguished experimentally by pharmacological inhibition of Erk. In this experiment, Erk activity would be temporarily impaired using a pharmacological inhibitor; then this inhibitor would be washed away, and the recovery of the system would be monitored. When the inhibitor is washed out, there is a before the system recovers and another wave is generated (Fig. 3C). The constant deposition model predicts a constant delay in wave onset determined by the time it takes for the activator to build up in the source region and trigger the positive feedback loop. By contrast, the oscillatory deposition model predicts a variable delay that depends on the phase of the deposition oscillation at the time of washout (Fig. 3D). Thus, experiments that analyze how new waves arise following perturbations of Erk activity could provide insights on the dynamic properties of the source. However, we note that this analysis may be complicated by potential additional feedback mechanisms between the rate of production of the activator and its downstream activity.

**Fig. 3.**
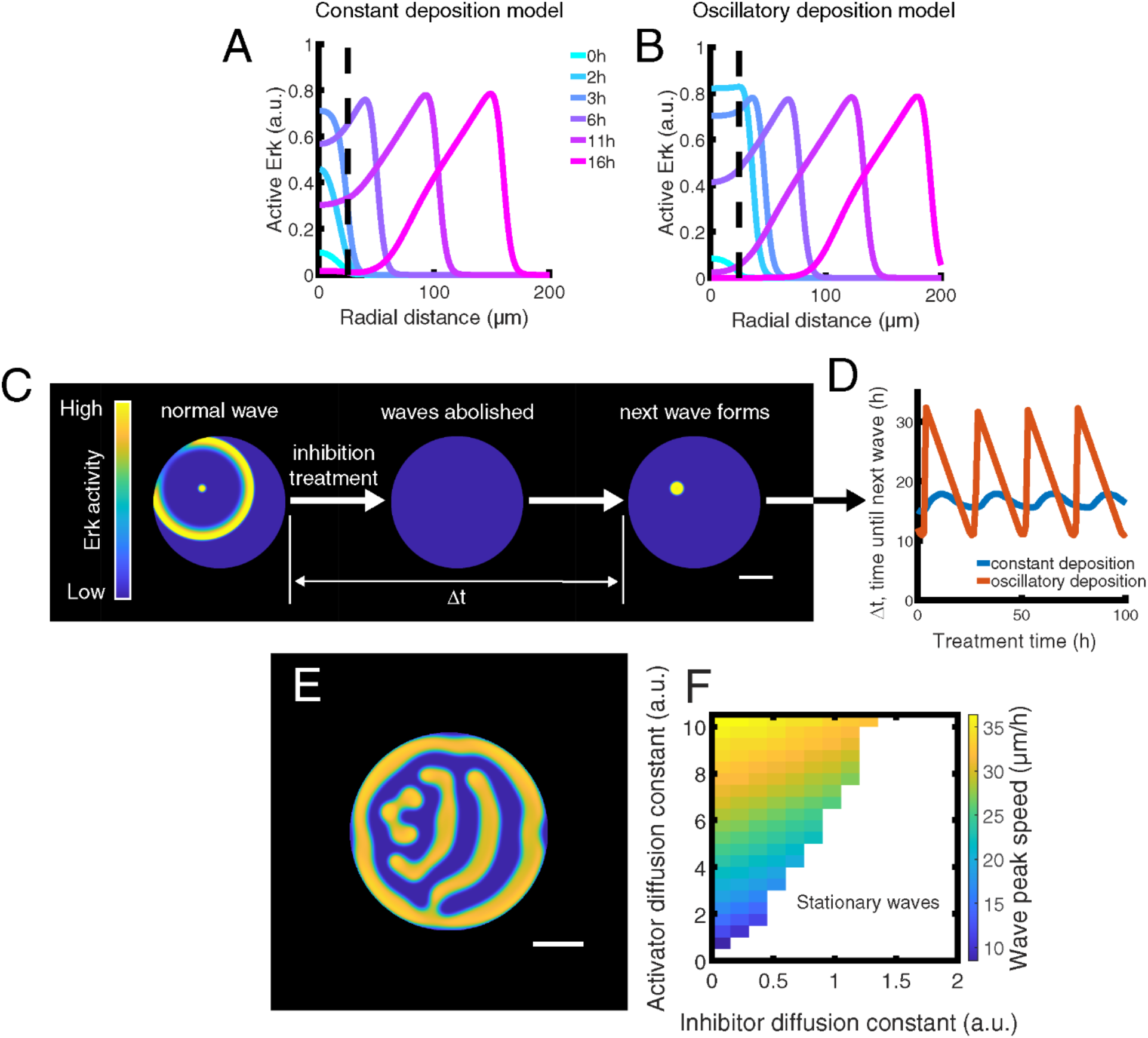
Variations in model type and assumptions. (A, B) Erk activity as a function of time in two models of wave generation at the source region. Dashed lines indicate the boundary of the source region. The standard model with a constant (A) or oscillatory (B) activator deposition in the source region. (C, D) *In silico* Erk inhibition and recovery experiment. Pharmacological inhibition is implemented in our model by setting *α*_1_ to 0 for the duration of the treatment. (E, F) Alternative wave propagation model in which the Erk inhibitor diffuses. (E) Example of stationary waves established after 200 h of simulation time. (F) Wave peak speed as a function of activator and inhibitor diffusion constants, relative to value of the activator diffusion constant in the standard simulation.

### Analysis of the effects of diffusion argue for waves as an effective mechanism to transfer information

The diffusivity of ligands *in vivo* is context-dependent, due to the complex microscopic details of ligand movement, and a wide range of different values have been observed for ligands of the Fgf family (7). Our experimental data on the speed of Erk waves (*υ* = 10 μm/h) and the typical timescale of Erk activation (τ = 3 h) allow us to estimate the effective diffusion constant of the signal propagating Erk waves as *D*~υ^2^τ~0.1 μm^2^/s (34). To further strengthen the evidence towards a small diffusion constant, we consider the possibility that an alternative model could generate the slow waves seen experimentally for larger values of the diffusion constant of the activator. In particular, we tested whether waves compatible with our experimental observations could be generated with faster activator diffusion if the Erk inhibitor could also diffuse among cells. However, we found that introducing inhibitor diffusion slows waves down only minorly, before waves are replaced by stationary patterns (Fig. 3E, F). We conclude that adding a diffusing inhibitor to our model cannot allow the activator diffusion constant to be much higher, given the experimentally observed wave speed.

To understand the physical implications of the low value of inferred diffusivity, we tested the ability of a diffusible morphogen (Fig. 4A) to propagate a signal across a domain of size similar to an adult zebrafish scale in a timescale like that of scale regeneration (34). To that end, we simulated the dynamics of an undegradable/unprocessed diffusible morphogen produced at the source. We found that for *D* ≈ 0.1μm^2^/s it takes on the order of weeks to achieve a significant morphogen concentration across the domain (Fig. 4B) while the wave mechanism in our model forms and propagates a wave across the same distance in just a couple days (Fig. 4C). Thus, we conclude that for values of the diffusion constant of the activator similar to the ones we estimated, morphogen gradients established by diffusion would be incapable of coordinating the cellular dynamics on the observed timescales.

**Fig. 4.**
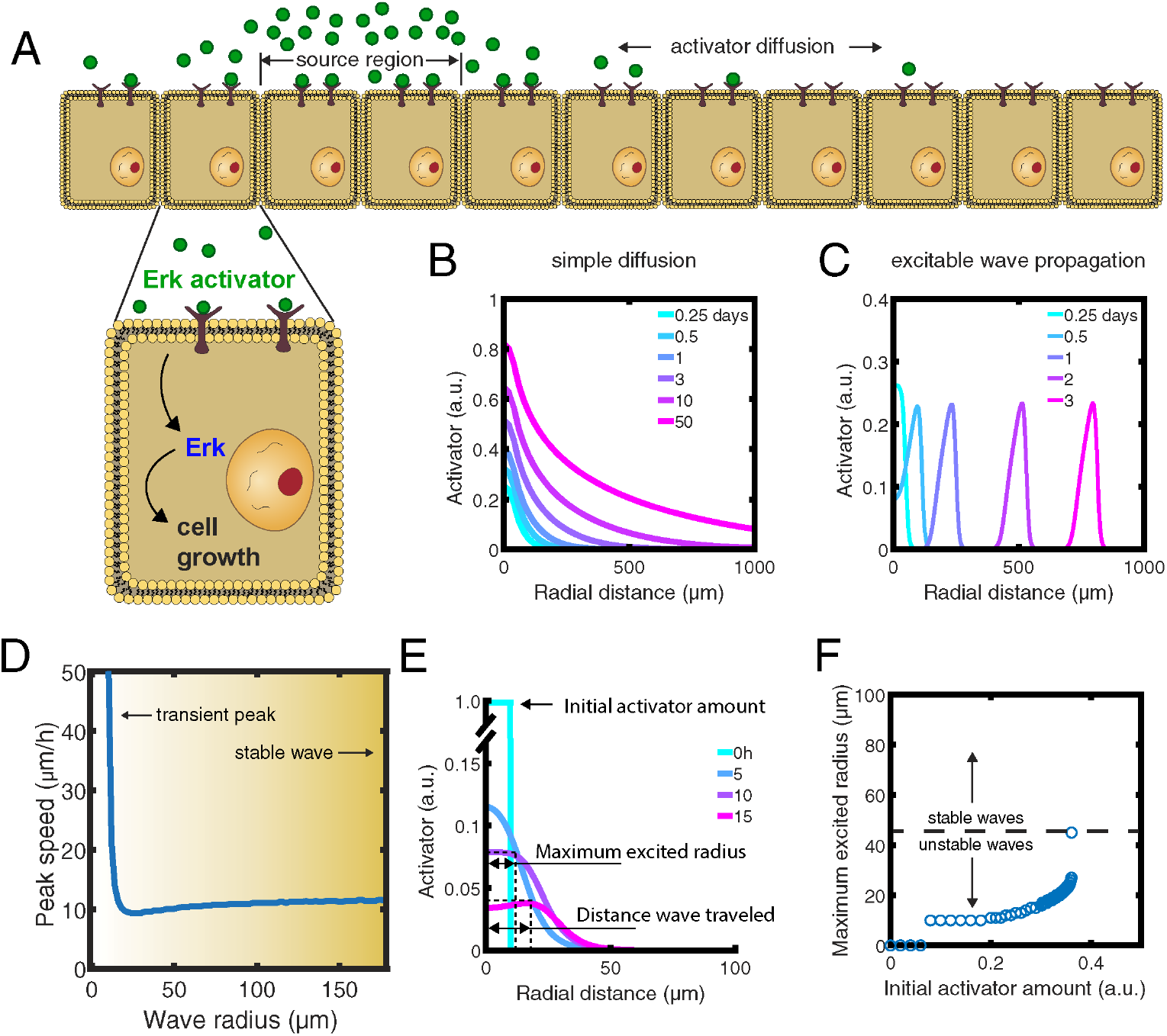
Consequences of activator diffusion constant and effects of curvature on wave propagation. (A) Schematics of activator gradient established by simple diffusion alone. (B, C) Activator gradient formation by simple diffusion (B) vs. excitable wave propagation (C). (D) Wave speed as a function of radial distance from the source. (E) Definitions of wave profile characteristics. (F) Radius of the excited region for different initial activator amounts. Initial activator concentrations higher than ~0.36 E_T_ induced an excited radius above 45 μm and triggered wave propagation.

### The effects of the geometry of the source on wave propagation

Wave geometry impacts the speed of excitable wave propagation (41). This relationship predicts that circular waves below a critical radius would not propagate. This radius is estimated to be *R_c_* ≈ *D/υ_p_* ≈ 45 μm in our system. However, in simulations, we can observe wave propagation even when the source region is smaller than the critical radius (Fig. 1D; Fig. 3A-B). We hypothesize that this is due to the fact that diffusion generates a larger profile of baseline activator concentration surrounding the source, thus an “effective source region”. The radius of this effective source region would be larger than that of the source by the order of magnitude of the activator diffusion length, 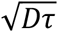. Compatibly with this idea, the initial spatially inhomogeneous profile of activator triggers the system excitation at different time, thus resulting in a high apparent speed of the Erk activity peak in and around the source (Fig. 4D). Thus, it is the size of this effective source region, and not the size of the source domain (i.e. where activator deposition *α*_2_ > 0), that determines whether a wave can propagate outward. To test this idea, we varied the amount of an initial pool of activator in the source region and allowed the system to evolve without additional activator production (Fig. 4E). The excited region was defined as the region in which the dynamic system passed the excitability threshold. Depending on the initial concentration of activator, different sizes of excited regions were established. Intriguingly, we found that when the excited region exceeded the predicted geometrical critical radius *R_c_* ≈ *D/υ_p_* ≈ 45 μm waves were generated, making the excited region as large as the scale domain (Fig. 4F). Thus, our results validate the idea that Erk activity waves can propagate only when their radius of curvature is above a certain threshold. These results further argue against a value of the diffusion constant much larger than *D* ≈ 0.1μm^2^/s, as for such values the radius of curvature for wave propagation would be significantly larger and occupy a large fraction, if not the entirety, of the scale tissue.

### A change in activator deposition can control wave frequency during regeneration

During regeneration of zebrafish scales *in vivo,* wave generation starts with a frequency of ~1 wave per day and slows down over time, until it stops (Fig. 5A-B). Wave frequency directly controls the rate at which scales grow over the timescale of regeneration (34). Our parameter analysis suggests that a change over time of the rate of deposition of the activator in the source region (*α*_2_) could explain this phenomenology (Fig. 2H). Alternatively, a change over time in the rate of degradation of the inhibitor *γ*_3_ could be responsible for reducing the frequency of wave generation (Fig. 2I). We combined experimental wave generation data with the numerical predictions on the effect of these parameters on wave period to infer the time dependency that either parameter would need to have to explain the experimentally observed change in frequency. We found that an approximately exponential decrease of the activator deposition rate at the source (*α*_2_) would be sufficient to explain this reduction (Fig. 5C). Thus, reducing the amount of activator delivered to the regenerating scale could be a mechanism to control the duration of regeneration and thus the size of the regenerating scale. This exponential reduction of activator deposition could be achieved, for example, if the activator at the source was deposited by a pool of dermal cells, each independently switching to a non-depositing state with a constant probability. Importantly, *α*_2_ controls only the dynamic of the source region, while the geometrical properties of the traveling excitable waves would be unaltered. Conversely, a reduction in γ_3_, which mainly controls the timescale of separation between Erk and the inhibitor, could explain the slowing of wave frequency but would result in a large increase in wave width (Fig. 5D). This can be understood by noting that for smaller γ_3_ the time required to extinguish Erk increases, and thus the width of the traveling Erk activity peak increases. Therefore, our results argue that the mechanism for controlling the duration of scale regeneration can be distinguished experimentally by monitoring waves throughout regeneration.

**Fig. 5.**
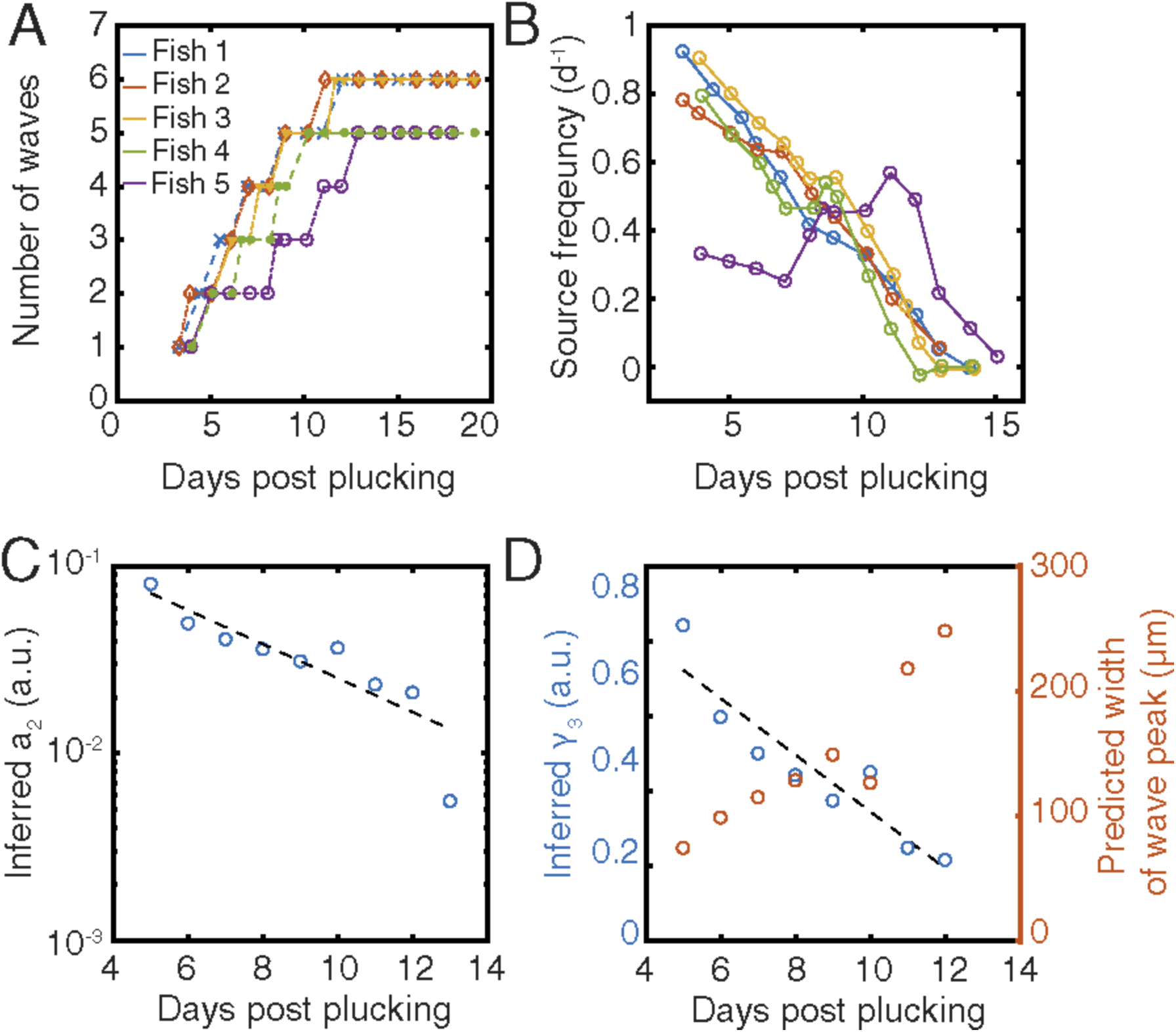
Effects of parameter changes on wave frequency. (A, B) Number of *in vivo* waves (A, panel adapted from (34)) and wave frequency (B) over the course of scale regeneration. (C) Inference of *α*_2_ from experimentally measured wave frequency. Dashed line: exponential fit *α*_2_ = *α*_2_(0)*e^−rt^* with *α*_2_(0) = (0.20 ± 0.05) and *r* = (0.21 ± 0.04)d^−1^. (D) Inference of *γ*_3_ and wave width over time. Dashed line: linear fit *γ*_3_ = *γ*_3_(0) + *rt* with *γ*_3_(0) = (1.1 ± 0.1) and *r* = (−3.2 ± 0.6) 10^−3^ h^−1^. Inferred parameter values are relative to the values in the standard simulation.

## Discussion

Waves can propagate stable signals across large tissues. We have recently reported that periodic waves of Erk activity, originating from a common source, control osteoblast regeneration in the zebrafish scale (34). Here we provide a mathematical model of Erk activity wave propagation in an excitable medium. At the core of this mechanism is an oscillatory source region, whose cellular nature and molecular regulation remain to be discovered. Once Erk activity is turned on in the source region, an excitable wave can travel across the rest of the tissue. Excitability requires positive and negative feedback mechanisms. Our analysis of the dependency of wave properties on model parameters supports the notion that the positive feedback between Erk and its activator controls the speed of the traveling wave. Conversely, the negative feedback between Erk and its inhibitor impacts the period of Erk excitable oscillations with little impact on wave speed. The periodic and repetitive nature of the wave is also controlled by the properties of the source region. Scenarios in which the ligand is continuously or periodically produced in the source region are both compatible with the observed repetitive nature of the waves. However, the two scenarios make different predictions on when a new wave would appear following inhibition of Fgf or Erk signaling. Testing this prediction experimentally could provide insights on the dynamical properties of the source.

Erk activity waves in scale regeneration present features that are different from the previously observed examples of Erk activity propagation. In scale regeneration, Erk activity propagates as a stable wave, with a constant amplitude. By contrast, in homeostatic mouse epidermis, Erk activity waves feature damped propagation up to ~50 μm and have a lifetime of ~30 minutes (25). In the epithelium of *Drosophila* tracheal placodes, waves propagate comparable distances over the course of about an hour (42). Similar to the case of scale regeneration, in wound healing assays, Erk activity waves can propagate from the edge of the wound for hundreds of microns, typically up to 200 μm. However, the remaining tissue displays a multitude of more disorganized waves that travel shorter distances (μ~ 100 μm) (43). Disorganized waves with origins influenced by noise are also seen in the activity of Cdk1 spanning the early *Drosophila* embryo to organize mitotic waves (44, 45). Thus, our experimental and theoretical work provide a rare example of a source region emitting repeated and coherent signals traveling across an entire growing and developing tissue. Clearly, future experiments should focus on discovering the nature and dynamic properties of the source region and how Erk oscillatory activity is controlled in such regions. In particular, it will be important to discover if the source region is created by interactions with other tissues, e.g. the vasculature or dermis, or if cells in the source region become specified by a self-organized mechanism which locks them in an oscillatory Erk regime, thus making them act as pacemakers.

The Erk waves in scale regeneration and in wound healing are characterized by different timescales. Most notably, the speed of Erk waves in regeneration is significantly slower than the speed observed in response to wound healing *in vitro* and in the skin epidermis (24, 25, 27). This slower speed is accompanied by an overall slower oscillation of Erk activity in individual cells. The duration of Erk activation is of the order of 10 hours in osteoblast regeneration compared to the 1 hour seen in the wound healing response. These slower dynamics have interesting implications for the possible mechanisms of wave propagation. In our experiments, the wavefront travels from one cell to the next in ~1-2 hours, a timescale compatible with a role of gene expression in wave propagation. Consistent with this idea, inhibition of *de novo* protein synthesis decreases the speed of wave propagation (34). Notably, a similar timescale/speed of propagation has been observed in the *Drosophila* eye imaginal discs, where the role of a transcriptional positive feedback is well-documented (46). In wound healing assays, however, the wavefront travels from one cell to the next on timescale of ~10 minutes, which suggests that mechanisms faster than transcriptional feedback are likely at play. Consistent with this scenario, a mechanism involving the cleavage of pro-activator molecules by cell contraction has been proposed (24). Thus, these observations suggest that different molecular processes can control Erk waves and that these processes might be selected for the relevant timescale of the biological process. Interestingly, the different mechanisms also employ different ligand families. Waves in regenerating osteoblasts are Fgf-dependent, whereas waves in wound healing are Egf-dependent.

The longer timescales of Erk oscillations also have implications on the possible mechanisms of the negative feedback that turns Erk activity off. Several negative feedback mechanisms exist that may regulate Erk. Some of these mechanisms involve phosphorylation by Erk, which can, for example, desensitize the pathway (47). These mechanisms would likely operate on rapid timescales and it is not obvious how they can be compatible with ~10 hours of Erk activation. However, mechanisms of negative feedback which involve the transcriptional activation of Erk inhibitors, namely proteins belonging to the Spry and Dusp families, are likely compatible with the timescales observed experimentally. In agreement with this, we previously reported that *spry* and *dusp* transcripts are enhanced in Erk active cells (34).

The geometric features of Erk waves have interesting implications on the role of diffusion. If the diffusion constant is high, molecules quickly move away from the source region, and as a result, the system would require a high activator production to surpass the threshold to begin a wave. Moreover, a high diffusion constant is predicted to lead to wide waves and a more uniform-like Erk activation profile (34). As a wave-like pattern of Erk activity is more conducive to regeneration than a wide activation of Erk (34), this further argues against a high value of the diffusion constant. Finally, the speed of a stable circular wavefront is negatively influenced by curvature-dependent effects on wave speed. These theoretical arguments suggest that the diffusion constant must be of the order 0.1 μm^2^/s, which implies that simple diffusion would take several weeks to form a gradient across the tissue. It will be important for future experiments to identify and study the receptors, ligand(s), and extracellular matrix determining the properties of this system. In conclusion, we speculate that waves provide two important advantages in the regulation of scale regeneration: rapid delivery of growth factor signaling and stereotypical pulses of fixed amplitude driving a similar amount of growth in all cells across the tissue. It is likely that excitable waves will emerge as a general principle of coordination of cell behaviors across large regenerating tissues.

## Materials and Methods

### Experimental data

Experimental data collection and analysis were described in (34).

### Numerical solutions of the model

The numerical solution of the presented mathematical model and analysis were performed with custom Matlab (Mathworks) 2019b software. The system of partial differential equations was simulated using the finite differences method (48). In the standard simulation, the simulation domain is a 1090 μm by 1090 μm square. Absorbing boundary conditions are set at the domain edge. A circle with a radius of 520 μm, centered in the simulation domain, is the scale region. Erk activity and the inhibitor concentration are set to zero outside the scale region. As a gap of at least 25 μm exists between the edge of the scale region and the domain edge and the activator degradation length is 7 μm, boundary conditions do not majorly affect system dynamics in the scale region. In the standard simulation, the activator source region is a small off-center circle with a radius of 40 μm. The standard time step is 0.01 h, and the spatial coordinates were discretized using a square lattice with grid size of 5 μm. The initial condition is that all chemical species are set to zero.

For parameter sensitivity analysis (Fig. 2; Fig. 5) and to reduce confounding effects due to wave curvature, we used a domain geometry that generates planar waves. Thus, in this case the simulations domain is a 10 μm by 1000 μm rectangle with a 10 μm by 50 μm rectangular source region at one end. Reflecting boundary conditions are used for the activator diffusion at the domain edges. Velocity was calculated by measuring the time required for the planar wave to travel 20 μm, starting from when a wave was 130 μm away from the source region. The analysis was limited to parameters that generate dynamics compatible to those observed *in vivo* (i.e. each source oscillation generates a travelling wave reaching the edge of the tissue). A source oscillation event was scored when active Erk in the source reached transiently a prominence above 0.2 (the “prominence” is the Erk active fraction at any given time minus the baseline Erk active fraction). A wave reaching the edge event was scored 500 μm from the source, when active Erk reached transiently a prominence above 0.2. Wave period was calculated by the duration between successive peaks in the excited region, 150 μm away from the source region. Wave width was calculated as Full Width Half Maximum when the wave peak was approximately 900 μm from the source region. Wave period, velocity, and width were calculated exclusively for parameter sets that led to the formation of a stable wave at every oscillation of the source region as is seen experimentally.

The oscillatory deposition model was similar to the standard simulation, but the activator deposition at the source *α*_2_ varied over time as a periodic Gaussian pulse:

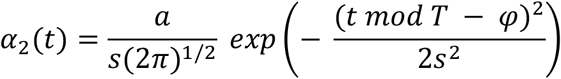

with amplitude *a*, standard deviation *s*, oscillation period *T*, and oscillation phase *φ*. *mod* indicates the modulus function, that is the remainder of the division of *t* by *T*. The oscillation period is set to 24.3 h to match the period of source oscillation in the constant deposition model. The oscillation phase is chosen as half the period. The standard deviation of the temporal oscillation was chosen to be 1 h to approximately match the activation and inactivation time of the wave. The gaussian peak amplitude was 2.8, calculated so that the total amount of activator deposited by the source over one oscillation period is approximately equal in the two models. To simulate treatment with an Erk inhibitor (Fig. 3D), *α*_1_ was set to 0 for 10 hours and then set back to its standard value. The time elapsed until the formation of the next wave was calculated as the time from the end of the treatment to that of the formation of a local maximum of Erk activity in the source region. In the model including inhibitor diffusion (Fig. 3F), inhibitor diffusion was treated in the same way as activator diffusion. To determine if stationary waves formed, simulation were performed for at least 200 h of simulated time.

To calculate the velocity of Erk activity waves as a function of radial distance (Fig. 4A) and to minimize the effect of the source radius on wave propagation, a 6 μm radius source region was used. To measure the minimum radius of the source excited region that can generate travelling waves, (Fig. 4D), the source activator production *α*_2_ was set to 0 and a certain concentration of activator was deposited in a circle of 10 μm radius at the onset of the simulation; each simulation was performed for 50 h of simulation time. At each time step, the excited region was calculated by testing for each grid point (grid size 1 μm) if it reached the region of the phase space in which an excitation occurs (in a simulation with null diffusion). The total excited region is the set of points that reached the excitation region at a certain moment during the simulation

In the model of activator simple diffusion (Fig. 4G), all model parameters were set to 0, except *α*_2_ and D.

In Fig. 5B, the experimental wave frequency was measured by smoothening the number of waves as a function of time and calculating a numerical derivative. Values of *α*_2_ and *γ*_3_ relative to the value in the standard simulation were inferred using the predictions of Fig. 2.

## Author Contributions

L.D.H., K.D.P., A.D., and S.D. designed research, L.D.H. and A.D. performed numerical simulations, L.D.H., A.D. and S.D. analyzed the numerical and experimental data, L.D.H., A.D., and S.D. contributed analytical tools; L.D.H., A.D., and S.D wrote the paper with comments from K.D.P.

## Acknowledgements

A.D. was supported by Early (P2ELP3_172293) and Advanced (P300PA_177838) Postdoc. Mobility fellowships from the Swiss National Science Foundation. This work was supported by an Innovation in Stem Cell Science Award from the Shipley Foundation, Inc. to S.D. and N.I.H. grant (R01-AR076342) to K.D.P. and S.D.

## Supplementary figures

**Fig. S1.**
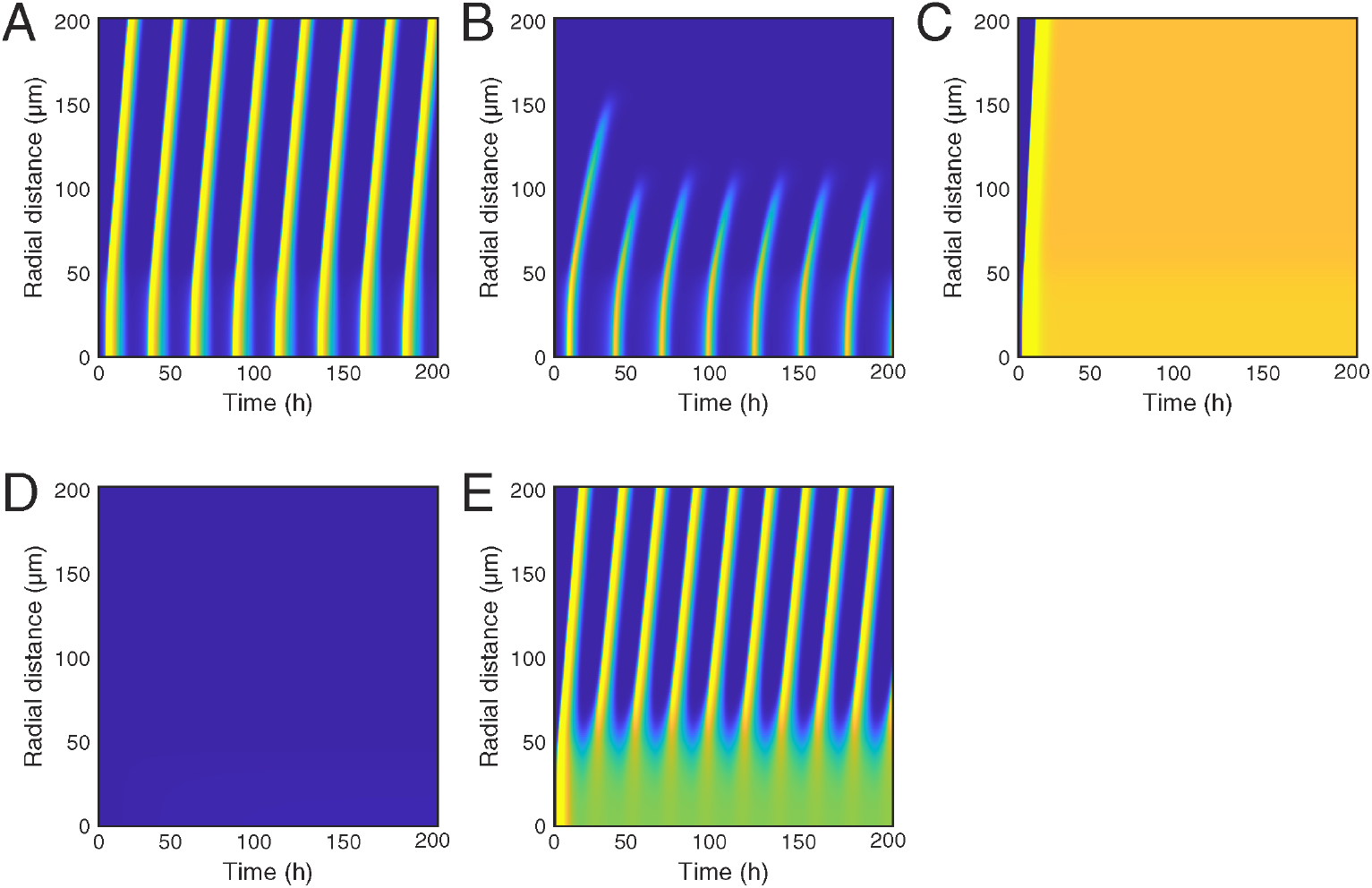
Representative kymographs of Erk activity dynamics in different regions of the parameter space. (A) Standard simulation. Periodic waves travel across the domain. (B) Damped waves, *α*_1_ = 0.63*α*_1_(*standard*). A fraction of waves does not reach the scale edge. (C) Bistable wave, *α*_1_ = 1.77*α*_1_(*standard*). A front of high Erk activity propagates across the simulation domain. (D) Waves are not generated, *α*_2_ = 0.15*α*_2_ (*standard*). (E) The source is locked in a high Erk activity state and travelling waves are periodically generated, *α*_2_ = 3*α*_2_ (*standard*).

